# Ultrahigh-throughput Absorbance Activated Droplet Sorting (UHT-AADS) for enzyme screening at kilohertz frequencies

**DOI:** 10.1101/2022.09.13.507731

**Authors:** Elliot J. Medcalf, Maximilian Gantz, Tomasz S. Kaminski, Florian Hollfelder

## Abstract

Droplet microfluidics is a valuable method to ‘beat the odds’ in high throughput screening campaigns such as directed evolution, where valuable hits are infrequent and large library sizes are required. Absorbance-based sorting expands the landscape of range of enzyme families that can be subjected to droplet screening by expanding possible assays beyond fluorescence detection. However, absorbance activated droplet sorting (AADS) is currently ∼10-fold slower than typical fluorescence activated droplet sorting (FADS), meaning that, in comparison, a larger portion of sequence space is inaccessible due to throughput constraints. Here we improve AADS to reach kHz sorting speeds in an order of magnitude increase over previous designs, with close-to-ideal sorting accuracy. This is achieved by a combination of (i) the use of refractive index matching oil that improves signal quality by removal of side scattering (increasing the sensitivity of absorbance measurements); (ii) a sorting algorithm capable of reaching 4 kHz with an Arduino Due; and (iii) a chip design that transmits product detection better into sorting decisions without false positives, namely a single-layered inlet to space droplets further apart and injections of ‘bias oil’ providing a fluidic barrier preventing droplets from entering the incorrect sorting channel. The updated ultrahigh-throughput absorbance activated droplet sorter (UHT-AADS) increases the effective sensitivity of absorbance measurements through better signal quality at a speed that matches the more established fluorescence-activated sorting devices.

**Table of Contents Graphic:** 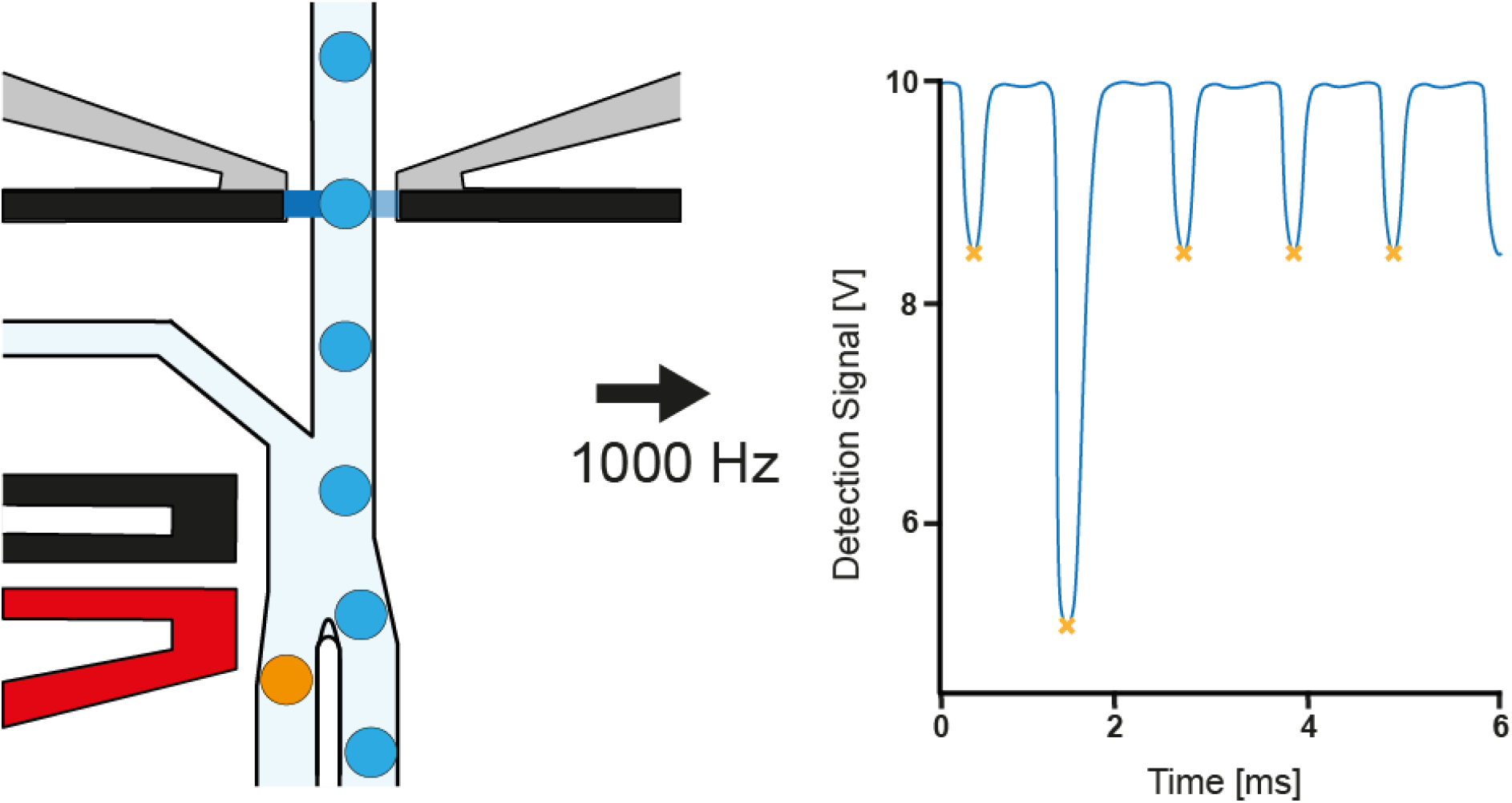

## Introduction

Functional screening of millions of different proteins in one experiment is possible at ultrahigh-throughput in droplet microfluidics, where water-in-oil emulsions co-compartmentalise genotype and phenotype. A selection is achieved by interrogating these pico- to nanoliter-sized ‘test tubes’ for an assay reaction that brings about an optically active primary or downstream product. Several successful directed evolution campaigns have been performed using droplet microfluidics to improve enzyme variants and develop non-natural catalytic properties^1–5^. Additionally, droplet microfluidics has also been used for functional metagenomics^6–8^, strain enrichment^9^, epistatic mapping^10^, and single-cell analysis^11^. Fluorescence assays are the most sensitive format for product detection: in fluorescence-activated droplet sorting (FADS), a few thousand molecules of a fluorophore in a droplet (corresponding to the nM range, e.g. fluorescein) can be detected with kHz rates^12,13^. However, the fluorogenic assays only cover a fraction of the reactions of interest, so alternative detection modes are required. Absorbance-activated cell sorting (AADS) provides practical means to cover chromogenic assays^14–18^, offering the opportunity to test a more comprehensive array of substrates.

Practically AADS is attractive: the setup is more straightforward and less expensive than FADS, as no lasers and photomultiplier tubes are needed. These factors also reduce safety requirements and mean that the microfluidic rig can be run on a lab bench instead of in a dedicated laser room. On the other hand, detection for AADS is not as sensitive as fluorescence detection (high µM vs nM detection limits, respectively)^12^. The AADS droplet sorter developed by Gielen *et al*. has been applied to the directed evolution of phenylalanine dehydrogenase^14,19^ and a glucose dehydrogenase^20^. Additional improvements to this initial design have been made: Zurek *et al*. overcame sensitivity limitations in droplet absorbance measurements by the growth of clonal variants in droplets^17^. Duncombe *et al*. introduced UVADS (UV-Vis Spectra Activated Droplet Sorter), which allows collection of whole spectra from 200 to 1050 nm, including a right angled turn at the detection interface to increase droplet path length for higher sensitivity^15^. However, the main limitation in all AADS designs remains the throughput at which droplets can be reliably sorted. The highest claimed sorting rate for absorbance detection is 300 Hz^14^ (although most enzymatic screening campaigns in our experience are generally performed at a lower throughput e.g. 100 Hz, due to a higher likelihood of incorrect droplet sorting at higher frequencies). This trade-off poses a limitation on the size of a library that can be screened in a practical timeframe and constrains the scope of the absorbance detection technology when trying to obtain rare variants in a screening campaign.

Compared to FADS, the second disadvantage of AADS is that absorbance is directly proportional to path length, meaning that droplets with a larger diameter and, therefore, larger volumes are needed. This, in turn, increases the amount of reagent required for each droplet and limits the throughput of sorting since a higher electric field is needed to sort larger droplets. If the electric field is too large, droplets tend to merge and/or become fragmented. A remedy to prevent merging is to provide a ‘Faraday moat’^21^ surrounding the channels upstream and downstream of the sorting junction; however, fragmentation remains an issue.

A third problem is of a practical nature, namely that the scattering caused by droplet edges as they pass the optic fibres places limits on the minimum droplet size and minimum substrate concentration that can be quantitatively detected. This effect becomes more pronounced for lower concentrations of the absorbing medium, as scattering obscures the true value of absorbance for lower molarities (i.e., the absorbance value is ‘hidden’ in between the edges).

A key challenge in directed evolution is the notion of ‘beating-the-odds’, such that the more variants screened, the higher are the chances of improving functional performance. Currently, FADS is at least ∼20-fold faster than AADS (1-2^1,3,6,7^ kHz vs 100 Hz^14^), implying a substantially reduced number of potential variants screened and leaving much sequence space unexplored. In the present work, we systematically explore various microfluidic, chemical and computational features of AADS and introduce improvements that collectively level its throughout with FADS, whilst still retaining high sensitivity, albeit at a higher volume than with a typical FADS set-up. Specifically, we address i) improved microfluidic design, ii) the use of added compounds for refractive index matching that are mixed with the spacing oil, and iii) development of new software for signal detection and automation of droplet sorting. Finally, the utility of our improved UHT-AADS (ultrahigh throughput AADS) was validated in an enrichment of phenylalanine dehydrogenase (PheDH) for the conversion of ʟ-phenylalanine (ʟ-Phe) to phenylpyruvate.

## Materials and Methods

### Chip Fabrication

Microfluidic chips were designed using AutoCAD 2021 (Autodesk) and fabricated using a high-resolution acetate mask (Micro Lithography Services Ltd.) in two layers. The detailed fabrication protocol is provided in the SI. Briefly, silicon wafers were spin-coated with SU-8 2050 photoresist (Microchem) and patterned on exposure to ultraviolet light. The second layer was aligned using an MJB-4 mask aligner (SUSS MicroTech). Microchannels were obtained from polydimethysiloxane (PDMS, Sylgard 184, Dow Corning) and were bonded to glass slides after surface plasma treatment. Channels were made hydrophobic by flushing of 1% trichloro(1H,1H,2H,2H-perfluorooctyl)silane (Sigma-Aldrich) in HFE-7500 (3M Novec). Device designs are deposited on our repository DropBase (https://openwetware.org/wiki/Dropbase:_UHT-AADS_Sorter) and their availability as CAD files immediately enable fabrication.

### Droplet Generation in a Flow Focusing Device

Droplets were made using flow focusing (height: 80 µm; width: 50 µm at the flow focusing junction) using HFE-7500 fluorinated oil and 2% 008-FluoroSurfactant (RAN biotechnologies). Flow rates were adjusted to the desired droplet volume e.g. for 75 pL the flow rates for oil and tartrazine solution were 40 µL/min and 10 µL/min, respectively. Gas-tight syringes (Hamilton Company) were operated with syringe pumps (Nemesys, Cetoni).

### Droplet Sorting

Droplets were sorted into the positive outlet channel by dielectrophoresis. Electric pulses with 1-1.2 kV at a frequency of 10 kHz were applied using on-chip 5 M NaCl electrodes. Light from the LED (455 nm, M455F3, ThorLabs) was passed between optical fibres (50 µm) at the detection point. A photodetector (PDA36A, Thorlabs) converted the reading to a voltage value from 0-10 V, which was passed into a DAQ (data acquisition) device (USB-6009, National Instruments) for real-time data visualisation using a custom LabVIEW script and into an analogue pin of an Arduino Due with a voltage divider to match the limit of tolerable voltage (3.3 V). On triggering from a positive droplet event, the Arduino Due sent a 3.3 V pulse to a pulse generator which delivered a 5 V pulse to a function generator to generate a 10 kHz AC square wave at 10-12 V. This was then amplified 100-fold with a voltage amplifier and coupled to the chip electrodes.

### Arduino Code and Data Analysis

The Arduino code was modified from Zurek *et al*.^17^ to additionally provide the ability to gate for droplet voltage and residence time. The code was written to maximise the speed of computation and run on an Arduino Due. Constant integer types were used, blocks of code were wrapped in functions, and the ‘micros’ function was used to allow continuous running of code with delays. A PicoScope 3000 Series (Pico Technology) oscilloscope was used to create an arbitrary waveform that resembled droplets passing the optical fibre. To determine the maximum frequency, the output of the Arduino Due was measured using the oscilloscope and the frequency of the arbitrary waveform was increased until the output of the Arduino Due did not trigger at the threshold value. New software is described in the SI and deposited on GitHub (https://github.com/fhlab/UHT-AADS).

### Enrichment of a Variant Expressing Phenylalanine Dehydrogenase

BL21(DE3) competent *E. coli* (New England Biolabs (NEB)) was transformed with a pASK-IBA63plus vector (IBA Lifesciences) harbouring a wild type PheDH strep tag fusion construct (Gielen *et al*.^14^) as positive control and a glycosidase as a negative control. The cells were grown to an OD600 of 0.4 – 0.6 in LB media and expression was induced by adding 200 ng/µl anhydrotetracycline. The proteins were expressed for 18-24h at 20°C and 200 rpm. A 1:100 dilution of positive to negative control in 20% Percoll (SigmaAldrich) was compartmentalized with substrate solution (4% cell lytic B (SigmaAldrich), 2 µl/ml r-lysozyme (Merck), 20 mM NAD (SigmaAldrich), 20 mM L-Phenylalanine, 15 mM WST-1 (NBS biologicals), 5 µg/ml mPMS and 2 mM tartrazine in 100 mM Glycine KOH buffer pH 8) in a flow focusing droplet generation device. Droplets were incubated overnight and the positive fraction was sorted. After sorting, 1H,1H,2H,2H-perfluorooctanol (Alfa Aesar) was added (1:1 ratio to oil), and the emulsion was broken by vortexing (1 min, full-speed). The oil phase was washed three times with 100 µl 2 ng/µl salmon sperm DNA (Invitrogen). The aqueous phase was purified and transformed into *E. coli* E. cloni 10G cells (Lucigen) by electroporation (1.8 kV). Cells were plated on LB agar supplemented with 100 µg/ml ampicillin and incubated overnight at 37 °C.

### Plasmid recovery and secondary assay

Plasmid DNA was extracted from the cells and transformed into *E. coli* Bl21(DE3) (NEB). Single colonies were picked and grown to saturation in 96 well plates. The cultures were diluted, grown to an OD_600_ of approximately 0.6 and protein expression was induced by addition of 0.2 mM IPFG. After expression for 18h (room temperature, shaking at 650 rpm), cells were pelleted (centrifugation at 4000 x g for 20 min) and resuspended and incubated in 200 µL lysis buffer (100 mM Glycine KOH buffer pH10, 4 mg/ml egg white lysozyme (Sigma) and 0.5 mg/mL polymyxin B (Sigma)) for 1h (room temperature, 650 rpm). The lysate was cleared by centrifugation. Enzyme assays were conducted in 96-well microplates with 10 mM L-Phenylalanine, 10 mM NAD in 100 mM Glycine KOH pH 10. Absorption at 375 nm was measured after incubation for 20 h.

For calculating enrichment factors equations previously used by Baret *et al*.^22^ and Zinchenko *et al*.^23^ were employed. Baret *et al*. define enrichment ɲ as the ratio of the ratio of positive to negative variants before (ɛ0) and after sorting (ɛ1). Zinchenko *et al*. define the ratio after sorting (ɛ1’) as a percentage of positive variants of the total pool of sorted variants and enrichment ɲ’ as ratio of ɛ1’ and ɛ0.

**Table 1:** Calculation of enrichment values

| $\epsilon_0^a$ | negative clones after sorting | positive clones after sorting | $\epsilon_1^b$ | $\eta^d$ | $\epsilon_1'^c$ | $\eta'^e$ |
| --- | --- | --- | --- | --- | --- | --- |
| 0.01 | 4 | 34 | 8.5 | 850 | 0.9 | 90 |

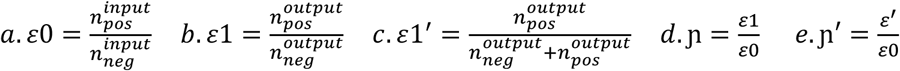

## Results and Discussion

### A device design for Increased droplet throughput

To achieve higher sorting frequencies, we first optimized the microfluidic sorting device through several iterations of design, fabrication, and testing of its performance. The design of a final device suitable for higher throughput is shown in Fig. 1A. The 2-layer chip features a bias oil inlet (see 3 in Fig.1A) for better spacing of reinjected droplets and also a gapped divider (see close-up in Fig. 1) at the sorting junction to minimise droplet fragmentation (adapted from Sciambi et al.^21^). The partial barrier (gapped divider) between the two outlets is designed so that the droplets do not break apart on impact and instead “hug” the sides of the barrier. This is important for larger-sized droplets at higher speed since a gentler impact ensures droplet stability. Large volumes inherently limit the sorting speed since as the volume increases, the electrophoretic forces required to move the droplet increase^14^, so smaller droplet volumes were employed in our study. Since absorbance is directly proportional to path length in line with the Beer-Lambert Law, decreasing droplet volumes reduces the diameter of the droplets and therefore leads to a lower absorbance that is harder to detect. There is, therefore, a trade-off between the droplet size and the maximum sorting speed. Also, as the droplet size increases, due to the shear forces acting on them, droplets are more likely to fragment due to the higher flow rate and higher electric field needed to direct droplets to the positive channel. The depth of the AADS device is determined by the width of the optical fibres. By using a shallower 50 µm-deep droplet injection chamber, smaller droplets can be evenly spaced, preventing two droplets from being injected at the same time and sorted incorrectly. These smaller droplets in a device with a depth of 100 µm are not squeezed, and therefore, ‘derailing’ them in the sorting junction becomes easier, so that higher sorting frequencies are achieved, which is also demonstrated in this study. The effect of a decreased droplet size and increased sorting speed is evaluated in the following experiments.

**Fig. 1.**
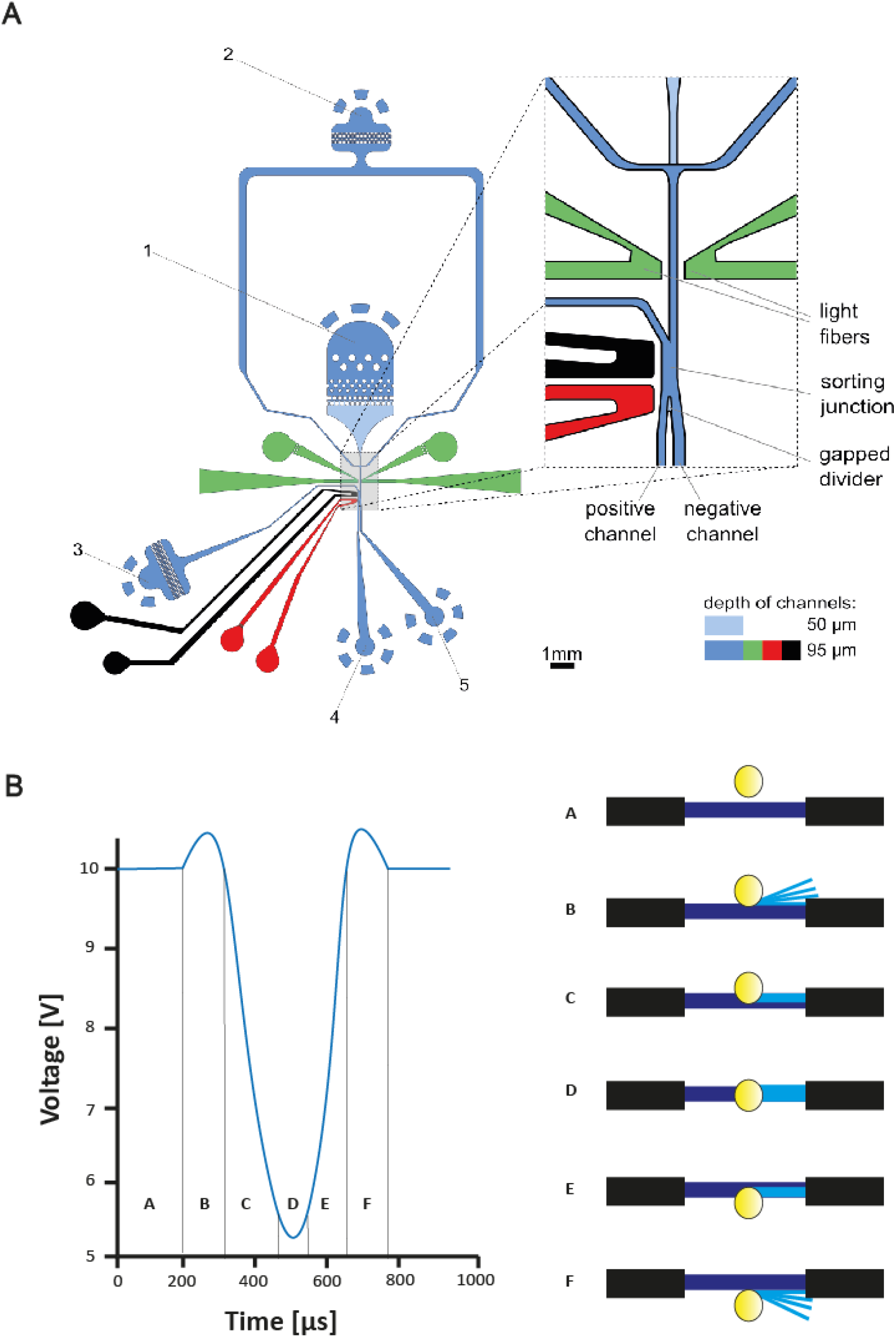
**(A)** A schematic of the UHT-AADS showing the single 50 µm layer channel (light blue) and a close-up design of the sorting junction showing the gapped divider at the bifurcation (light blue). The gapped divider is on the second 50 µm layer. 1) input channel for droplets, 2) input channel for spacing oil, 3) input channel for bias oil, 4) positive channel outlet, 5) negative channel outlet, 6). The ground electrode is coloured black and the positive electrode is coloured red. Adding a single-layer droplet injection chamber allows for even spacing of the 75 pL droplets. The bias oil channel acts as a barrier to droplets from entering the negative chamber unless acted on by the dielectrophoretic force. **(B)** A diagram showing a typical droplet trace as it passes the optical fibres. At position A, light transmission is at the maximum value indicated by the baseline value set. At B, the droplet edge causes refraction at the water-oil interface producing a ‘shoulder’ corresponding to the higher amount of light collected by the fibre. At C, the droplet is moving towards the centre of the optical fibres. At D, the droplet is in the centre of the optic fibre, and the true peak value of absorbance (minimum amount of light) is given in the droplet trace. At E, voltage increases as the droplet moves away from the optical fibre centre. At F, there is the effect of the other droplet edge causing another peak (or a ‘shoulder’) of collected light.

### Software and Electronics

Higher throughput of optimized microfluidic devices required implementation of improvements in the software that was used for recording the signal and triggering sorting events.

#### (i) Sorting algorithms

As throughput and speed are improved through the device design, the speed of computation for sorting decisions is challenged. It is important to consider the algorithms for droplet sorting since the maximum speed of sorting cannot be higher than the time needed for computational decision making (determined by the maximal computation speed). Previously the algorithm used in Gielen *et al*.^14^ was limited to triggering by a simple threshold value which made setting a stable gate more difficult. The use of a delay function meant that sorting would be limited in speed if a gating step was added. The open-source Arduino software developed in Gielen *et al*.^14^ was therefore redesigned (incorporating gating code modifications from Zurek *et*.*al*.^17^) to include gating of droplet residence times and voltages and the possibility to sort for ‘negative’ droplets (i.e. droplets with a lower absorbance than the rest of the population) was added. These improvements allow much more precise control over which droplet events are sorted, such that anomalous events (e.g. passing of an air bubble which usually triggered the electrodes due to a decrease in voltage), are identified and deselected due to gating. Likewise merged droplets and other artefactual events (e.g. dust passing the optic fibres) can also be excluded. Additionally, better control over sorting thresholds increases applicability of the setup to directed evolution campaigns where sorting gates need to be strictly controlled to avoid sorting of inactive variants.

#### (ii) Code

To optimise the speed of the code, static variables were converted to a constant integer type, the micros function was used to allow delays without blocking the program, and blocks of code were wrapped in functions. The code has also been designed to run at speeds in line with the increase in droplet throughput. In simulated tests of purely electronic response rates, the maximum throughput of the new algorithm is 4 kHz using an Arduino Due device (see the script in the SI). Custom Python scripts were created to process the raw data from the photodetector (see scripts in SI). The SciPy package^24^, alongside others, allowed peak detection of droplets, determined the baseline value, and further explored the data.

#### (iii) Electronics

The sorting electronics originally designed by Gielen *et al*.^14^ included an Arduino Due, which sends a pulse at 5V to a pulse generator which then triggers a Function Generator linked to a Voltage Amplifier to pulse at 10 kHz at between 6-10 V. This avoided the use of a FPGA (Field Programmable Gate Array) to measure input data and trigger pulses. Even though FPGAs enable parallel processing, and their hardware is programmable, together resulting in computation speeds that are higher than microcontrollers, expertise in low-level programming such as Verilog or VHDL is required, limiting both the accessibility and reconfigurability of these devices for the microfluidics community^25^. By contrast, our Arduino-type microcontroller can be programmed using higher-level languages (decreasing complexity and increasing accessibility), are open-source with significant community support, incur low costs and their code can be modified more easily compared to FPGAs. Given that processing complexities for droplet sorting are minimal, and with the increasing processing speed of microcontrollers, we argue that using an FPGA for most droplet sorting applications is redundant since the extra complexity is not needed.

### Baseline offset and refractive index matching of the oil phase create smoother voltage peak shapes

We additionally focused on implementing a new strategy for the removal of unwanted scattering signals coming from light deflected by the droplet interfaces. The determination of the true absorbance value becomes much more challenging due to the presence of the spikes caused by the droplet edges. This makes it algorithmically harder to assign a true absorbance value correctly (see Fig. 3, 150 pL negative control) and takes additional time to compute the correct value, complicating real-time signal processing. Specifically, readouts of the droplet absorbance by monitoring voltage over time (Fig. 2B) showed scattering at the droplets’ edges at the oil-water interface of droplets as they pass through the optical fibre detection area (Fig. 1B). This effect is due to refraction and causes unwanted spikes in the voltage data, so that the true absorbance value of the droplet is only visible in between the spikes. So far, the problem of spikes was bypassed by the addition of a dye offsetting the signal (Gielen *et al*.^14^). At the saturation limit of the photodetector droplets with low molarity of absorbing dye can have an absorbance value greater than that of the saturation limit, lifting the true value above the saturation limit (and therefore not quantitative). To minimize this effect, an offset of tartrazine can be added to the droplet contents, resulting in a deliberate absorbance increase to a level below the saturation limit. However, doing so reduced the dynamic range, as the detection range was lowered.

**Fig. 2.**
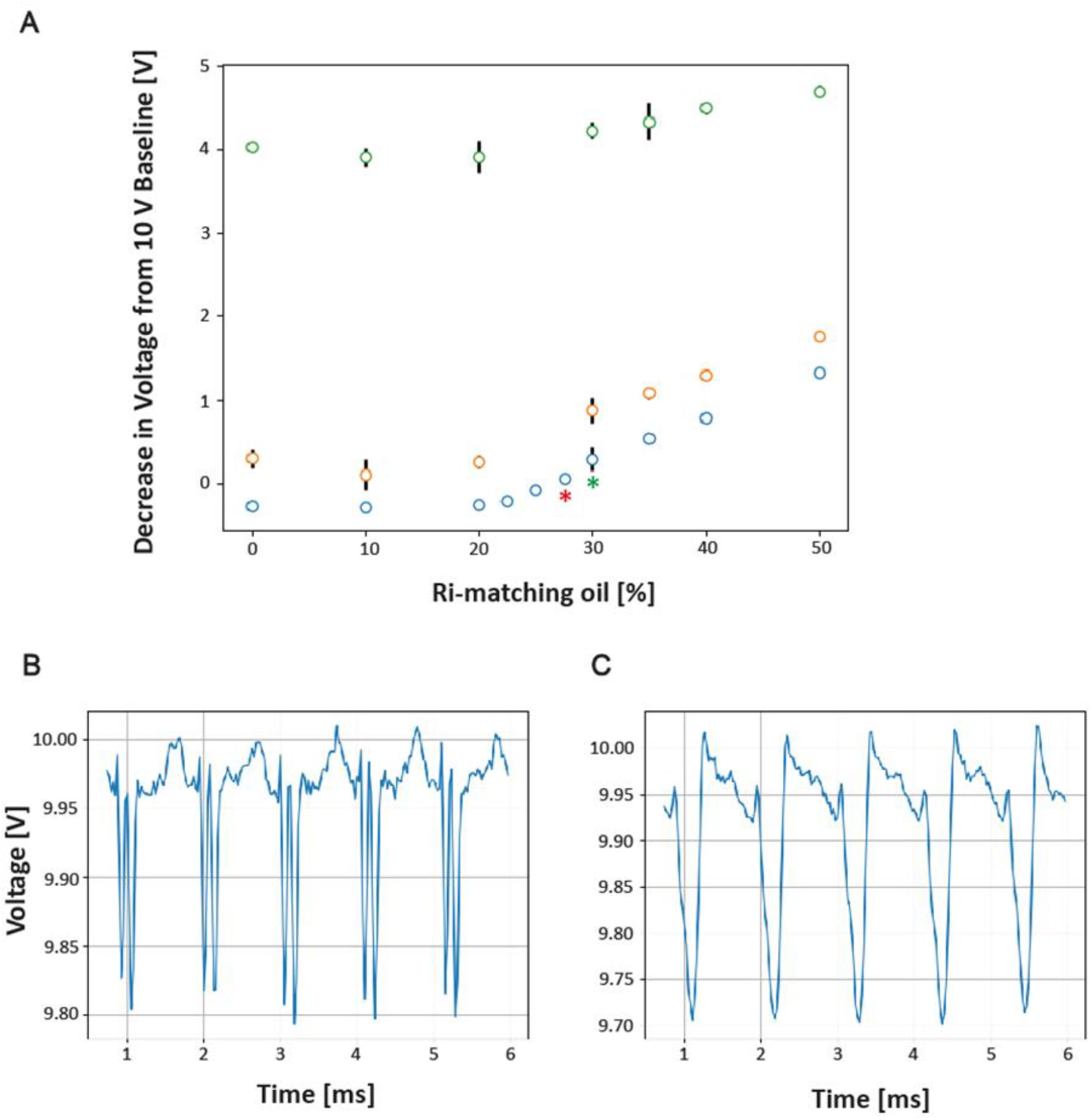
**(A)** The decrease in voltage from the 10 V baseline shows that as the RI-matching oil percentage increases, so does the decrease in voltage from the baseline value. The gradient decreases as the concentration of tartrazine increases, showing the 0 mM (blue) and 0.5 mM (orange) having steeper gradients than 5 mM (green). The voltage at which the RI-matching oil has no bias effect is shown at ∼25%. Each point is data from 20 seconds of droplet voltage recordings at 1000 Hz. Error bars are the standard deviation of the 20,000 droplet peaks extracted from the data. The red asterisk indicates the droplet trace for B, and the green asterisk indicates the droplet trace for C. **(B)** 6 ms droplet trace for 0 mM tartrazine (in water) using 27.5% RI-matching oil. Droplet values are hidden between the droplet’s edges, making it difficult to discern the real peak of the droplet. (**C**) 6 ms droplet trace for the negative control (0 mM tartrazine) using 30% RI-matching oil. Droplet peaks are clearly distinguished from the baseline, and droplet edges have been removed.

**Fig 3.**
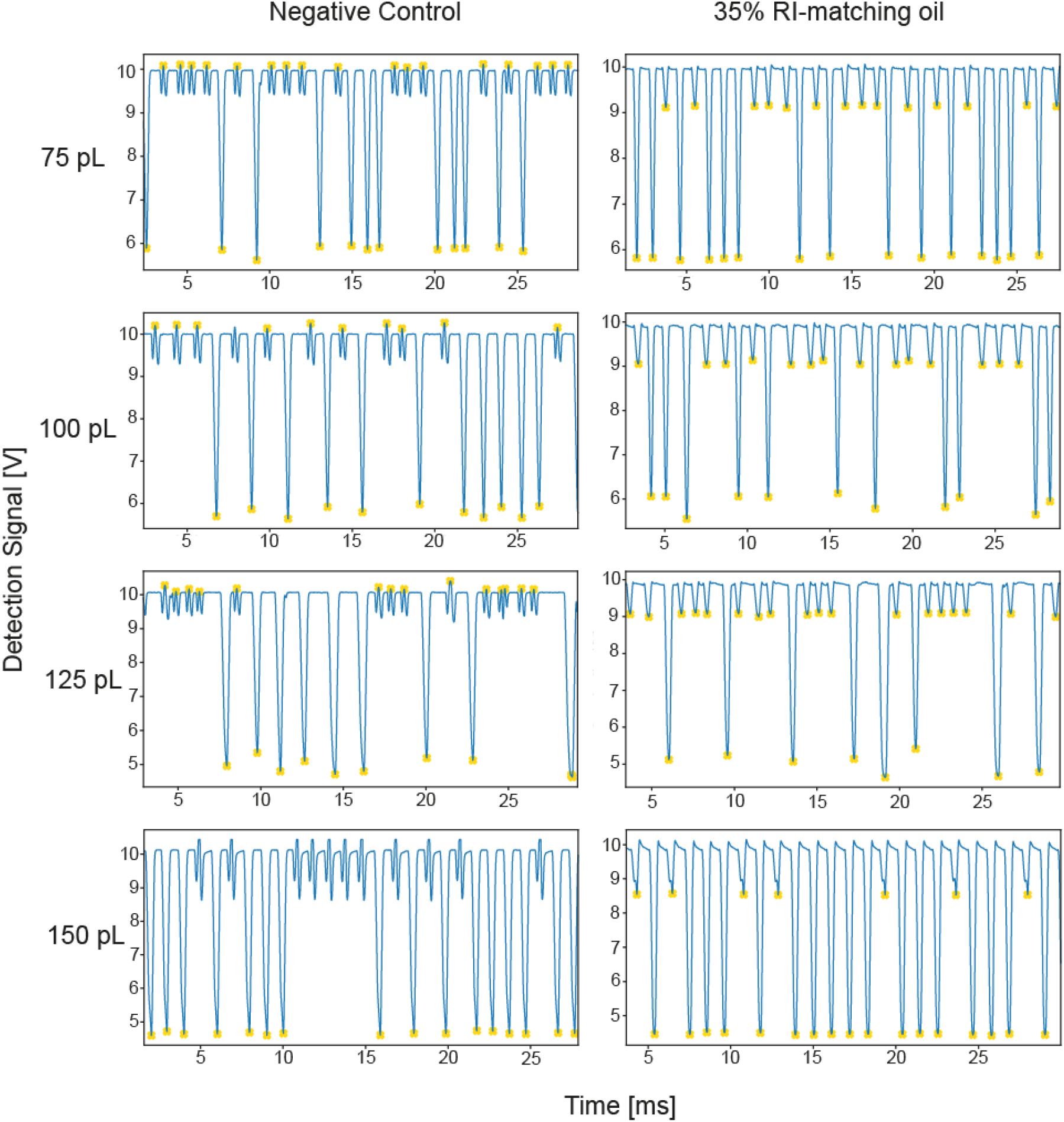
The effect of RI-matching oil on the droplet traces (V) at different droplet sizes with two droplet populations, 0.5 mM tartrazine (peaks at higher voltage) and 5 mM tartrazine (peaks at lower voltage) at approximately 1 kHz frequency of droplet measurement. The negative control is without 1,3-bis(trifluoromethyl)-5-bromobenzene and shows significant droplet edges for all droplet volumes. No droplet edges are seen when using 35% RI-matching oil and peaks are clearly distinguishable. Droplets with a volume of 75 pL are distinguishable at 0.5 mM. The yellow asterisks show droplet peaks identified using a custom peak detection algorithm (SI). The algorithm cannot distinguish the droplet values for the 150 pL negative control trace due to broad peaks leading to an unidentifiable maximum.

In an alternative approach to mitigating the issue of scattering at droplet edges, 1,3-bis(trifluoromethyl)-5-bromobenzene was added to the fluorocarbon HFE-7500 oil (Salmon *et al*.^26^), as an example of a refractive index (RI) matching compound. This RI-matching oil has high miscibility with oil and does not interfere with droplet re-injection and spacing in the sorting junction. Several additional practical considerations determined how such a mixture best removed the droplet edges in the signal, provided a wide dynamic range, and allowed empty droplets to be confidently identified. The RI-matching oil had an additional effect on the measured absorbance values. The choice of a suitable percentage of RI-matching agent is determined first by trying to minimise the scattering effect, and secondly, to have a suitable dynamic range between high and low concentrations of absorbing compound (e.g. tartrazine). Figure 2A shows how increasing the percentage of 1,3-bis(trifluoromethyl)-5-bromobenzene leads to an increase in voltage although the absorbance of tartrazine, here used as a model absorbing compound, does not change. Indeed, the gradient is increased for 0.5 mM and 0 mM tartrazine concentrations beyond 30% RI-matching oil. This indicates that increasing the concentration of RI-matching oil decreases the assay’s dynamic range since there is a lower range between e.g. 5 mM tartrazine and 0.5 mM tartrazine at higher concentrations of RI-matching oil. Fig. 2B shows that for a RI-matching oil concentration of 27.5% there are still significant droplet edges, and the true peak lies in between. However, it is necessary to detect empty droplets for frequency and other measurements (e.g. gating in directed evolution screening campaigns), so a suitable concentration of RI-matching oil should also give enough of a voltage signal to allow a signal to be detectable for these droplets (albeit without droplet edges). Values with a RI-matching oil percentage of greater than 30% do not show droplet edges in the trace and therefore their true value is hard to deconvolute (Fig 2C). Overall, the value chosen is 35% 1,3-bis(trifluoromethyl)-5-bromobenzene since this represents the best compromise between removing droplet edges, providing a high dynamic range, and allowing empty droplets to be identified. Removing edges significantly minimises the sorting algorithm’s complexity (since it is difficult to deconvolute the true value from traces with edges) and therefore increases computation speed.

The addition of an RI-matching compound to the spacing oil also required a change in the design of the microfluidic sorting junction. Since the RI-matching oil reduces emulsion stability and might cause wetting of droplets to walls of channels or tubing, we decided to apply a strategy of transferring positively sorted droplets back to the oil without RI. Therefore, a bias oil channel (through which standard HFE-7500 oil with 0.5% 008-FluoroSurfactant, i.e. with no RI-matching compound) was added at the sorting junction and during sorting both oils are a laminar flow regime. This standard oil, therefore, poses a barrier to droplets from entering the positive outlet. Only when the electric pulse applied to the droplets from the sorting electrode exceeds the inertial force driving the droplet into the negative outlet does the droplet enter the positive outlet (see Fig. 1A for the diagram).

### Effect of droplet size on absorbance

Fig. 3 compares the effect of different volumes for two droplet populations (0.5 mM tartrazine and 5 mM tartrazine) measured at 1 kHz with and without 35% RI-matching added to the spacing oil. As the droplet volume decreases for both the negative control and 35% RI-matching oil, the detected voltage also decreases, potentially due to the decreased path length of the droplet resulting in a lower absorbance value. As shown in Fig. 3, the droplet traces for 75 pL droplets with RI-matching oil show clear peaks with no shoulders. However, without RI-matching oil, the scattering effect causes the true droplet value to be above the baseline and is masked by the higher absorbance of the spacing oil. The measurement, therefore, is not quantitative as the droplet values are past the saturation limit.

As previously stated, droplets with a smaller volume are easier to sort at high frequencies due to less force needed to move them effectively. Our results proved that modified microfluidic design and adding RI-matching oil allow for detection and sorting of droplets with a volume of 75 pL. This is a significant improvement compared to larger volumes of droplets in previous studies (e.g. 180 pL in Gielen *et al*.^14^)

### Sensitivity of detection

Recalibration of the sensitivity is needed in reference to Gielen *et al*.^14^ since the absorbance values are affected by the smaller path length. The range of the calibration plot was generated by converting to the mean absorbance for 2000 droplets (Fig. 4). As can be seen, there is a positive linear trend as the concentration of tartrazine increases from 100 µM to 5 mM, as expected. Below this, the trend does not continue showing that the device’s sensitivity is around 100 µM. Since smaller droplets are used to sort at higher frequencies, the path length is decreased, and in accordance with the Beer-Lambert law, the dynamic range is decreased. The advantages however with smaller droplets, as previously mentioned, are a higher throughput, a higher local concentration of potential enzyme, and a smaller amount of chemical solutions needed. As can be seen in Fig. 4, the standard deviation of the absorbance values is small, indicating that the precision of detection is high.

**Fig 4.**
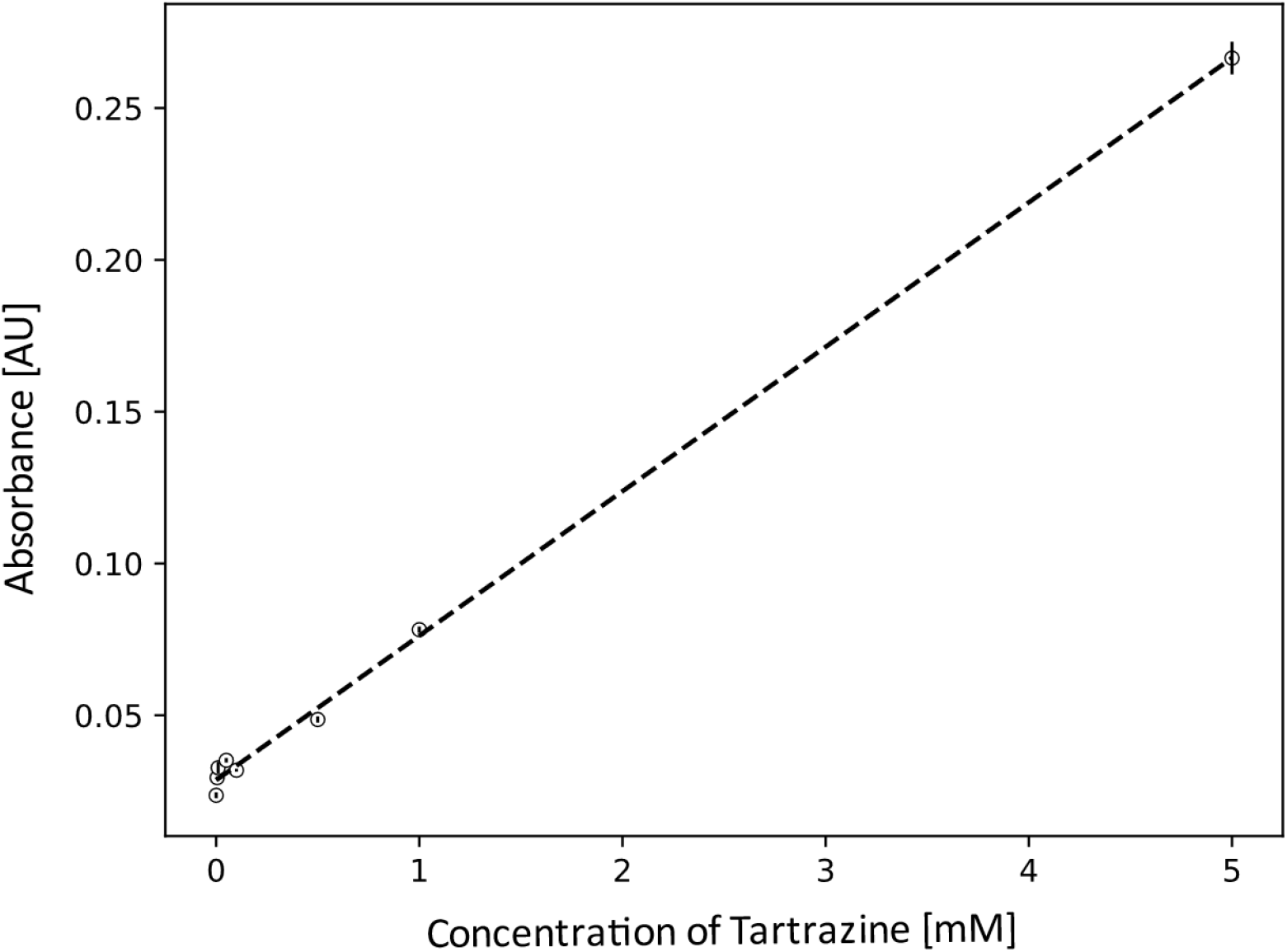
Calibration plot showing the linear relationship of absorbance to the concentration of tartrazine (mM) which is used as a model absorbance compound. A residence time gate was applied to remove artefacts such as air bubbles or merged droplets. Two milliseconds of data (around 2000 droplets with a volume of 75 pL) were measured, and the peak values were determined. Absorbance was calculated using the equation *A* = −*log*10(*V*_*n*_/*V*_*0*_), where V_n_ is the voltage at a given concentration and V_0_ is the voltage for pure RI-matching oil. 35% RI-matching oil was used for spacing of droplets. Error bars are the standard deviation of the ∼2000 droplets.

### Validation of sorting efficiency at different droplet frequencies

To test whether the improvements described in the previous paragraphs do not only allow fast sorting, but also high efficiency in the sorting outcome, we carried out selections (between two 75 pL droplet populations, 0.5 and 5 mM of tartrazine, respectively) at different frequencies (1, 1.5, and 2kHz). Flow rates were adjusted to allow different droplet frequencies and the ratio of flow rates was kept the same between frequencies (1:5:10 ratio for droplet: bias oil: spacing oil, respectively). Droplets were counted as being correctly sorted only if the droplet triggering the sorting event was sorted alone, without any fragments from other droplets before or following behind. Fragmentation of droplets occurs as the frequency increases due to the insufficient force required to move the droplet into the positive outlet, and the droplet collides with and breaks at the barrier. Additionally, as the electric field increases this also leads to droplet fragmentation. This is also more pronounced because of the different oil composition at the sorting junction. Figure Fig. 5C shows an example of a sorting event recorded at a high frequency of 2 kHz, suggesting that the force required to move the droplet into the positive outlet chamber is not sufficiently strong: the droplet collides with the centre of the barrier, splitting it into two daughter droplets. A potential remedy would be to use a greater electric field to move the droplet into the positive outlet more strongly before it collides with the barrier. However, this may also inadvertently cause the droplet to break up due to increased instability at the droplet surface.

**Fig. 5.**
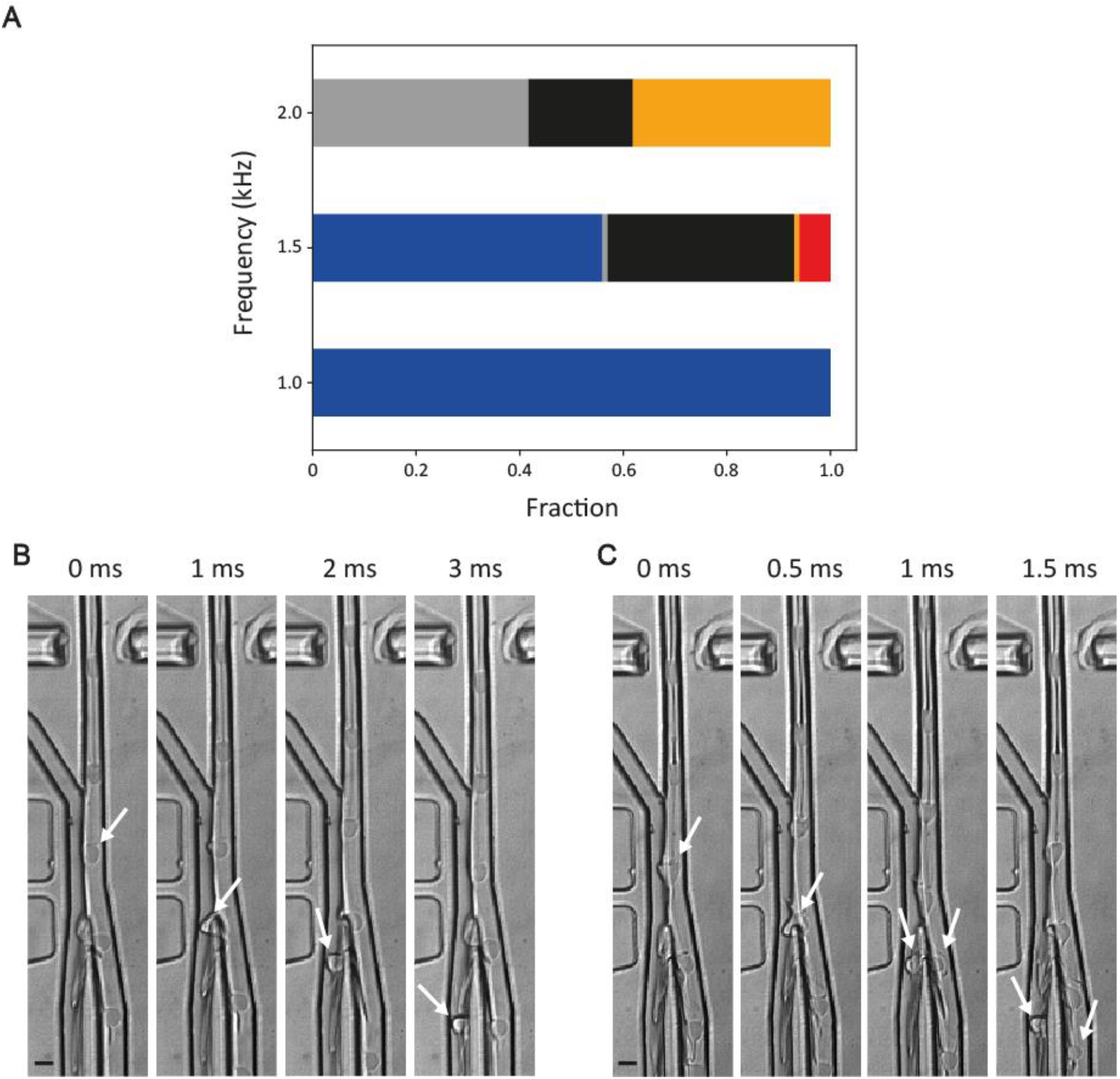
**(A)** Bar chart showing the fraction of correctly sorted droplets (blue), partial false negatives (grey), false negatives (black), partial false positives (orange), and false negatives (red) at 1, 1.5 and 2kHz. Partial false negatives are droplets that are false negatives that split at the junction. Partial false positives are false positives that split at the junction. A high-speed camera was triggered at every sorting event, and visual inspection in ImageJ was used to determine if the droplet moved into the correct outlet. Droplets were counted as correctly sorted only if the droplet that triggered the event was sorted alone. The events examined were ∼100. **(B)** Snapshots of droplets at 1000 Hz when the sorting electrode is triggered based upon the correct absorbance value. The droplet that is correctly sorted is shown with the white arrows. **(C)** An example of a droplet fragmenting at 2000 Hz due to collision with the central barrier. Arrows show the droplet splitting into two sub volumes.

Finally, since the bias oil and flow rate of the droplets are entering the junction at different rates, this poses an additional barrier to pulling the droplet into the positive channel. As the droplet is pulled over to the positive elution channel part of it is still being pulled into the negative outlet at a higher flow rate. This stress on the droplet causes the droplet to break apart at the barrier.

When probing higher droplet frequencies, the forces needed to pull the droplet into the positive outlet chamber must become larger. Accordingly, sorting efficiency decreases as the droplet frequency increases (Fig. 5A), and the number of droplets that become fragmented also increases (partial false negatives and false positives). All in all, a compromise between the high voltage needed to pull the droplet into position and a low enough voltage that does not cause droplet fragmentation are needed to achieve a balance between droplet and bias oil flow rate to ensure minimisation of false positives and enough giveaway to sort positive droplets correctly.

In the light of these issues the success of sorting was evaluated by analysing video traces (with ∼100 droplets after sorting) with an algorithm detecting the absorbance in droplets passing the detector. Sorting was carried out at one kilohertz (100% efficiency for 100 videos analysed), a 10-fold improvement of the apparatus used by Gielen *et al*.^14^.

### Validation of the sorting accuracy by enrichment of active phenylalanine dehydrogenase variants in a single cell lysate screening experiment at 1 kHz

An enrichment experiment was performed in a droplet-based colorimetric cell lysate assay for phenylalanine dehydrogenase (PheDH) activity using the workflow established here. WT PheDH and an enzyme lacking PheDH activity (glycosidase-expressing as a negative control) were mixed in a ratio of 1:100. We then screened for droplets containing the wt PheDH at the improved sorting rate of at 1 kHz. DNA was recovered by transformation into E. coli and the output analysed in a secondary photometric assay to evaluate the efficiency of sorting. A small sample was analysed spectrophotometrically (Fig. 6C) and 34 of 38 clones tested showed PheDH activity (within two standard deviations from the mean of the positive), giving a true positive rate of 89%. This translates into a 90-fold enrichment calculated according to Zinchenko *et al*.^23^ and an 850-fold improvement according to Baret *et al*.^22^, suggesting that the enrichments are suitable for functional selections.

**Fig. 6.**
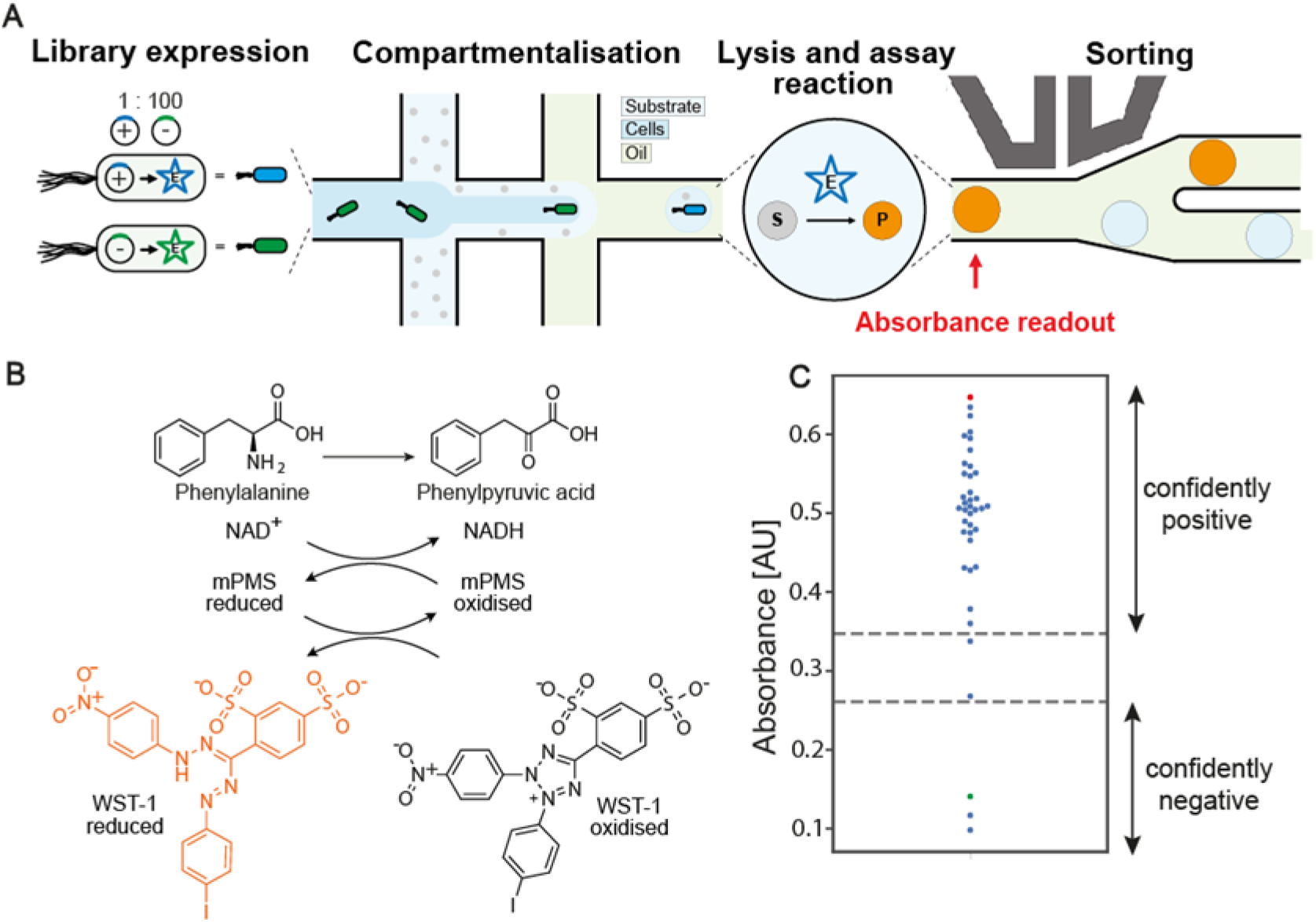
**(A)** Illustration of a screening workflow, showing *E. coli* expression of a library, droplet generation for single cell encapsulation of library members and sorting for active phenylalanine dehydrogenase (PheDH) activity. **(B)** The coupled reaction produces reduced WST-1 with absorbance at 455 nm. **(C)** Swarm plot of the mean absorbance at 340 nm for 38 colonies (blue, n=3 for each) picked after transformation of DNA collected from the positive outlet of the AADS device sorting at 1 kHz. The red dot shows the mean (n=9) of the positive control (pASK expressing wild-type PheDH), and the green dot shows the mean (n=9) of the negative control (pASK expressing glycosidase). The confidently positive clones are classified as two standard deviations from the mean of the positive control and respectively for the negative control (dotted lines).

## Conclusion

Absorbance-activated droplet sorting is still less frequently used for protein engineering^14,17^ compared to the more established FADS, despite their complementarity and the possibility of screening a larger and different pool of potential reactions. This work enables AADS to catch up in its performance; the 1 kHz sorting rate (with 99% efficiency) is equivalent to FADS campaigns. Based on this 10-fold improvement of throughput (compared to 100 Hz reported in biological experiments previously^14^) larger libraries can be screened in the same amount of time, increasing the chances of success by making it more likely to identify rare functional variants as larger fractions of sequence space are interrogated.

Other drawbacks of AADS remain: for FADS, the minimum product concentration is >2 nM (corresponding to >2500 product molecules per droplet), whereas for AADS the sensitivity is lower at >7.5 µM (with >10^9^ molecules per droplet)^13^. In fact the sensitivity of the set-up described in this work is slightly lower than that introduced by Gielen *et al*.^14^ due to the path length reduction. However, the local enzyme and product concentrations are larger in smaller droplets (2.4-fold in 75 vs 180 pL droplets), partially compensating for the reduced sensitivity. In-droplet growth of enzyme-expressing *E. coli* cells (after single cell encapsulation) has been shown to increase the amount of enzyme in an assay, which provides means to boost the product signal (while also reducing phenotypic variation)^17^. Increasing the sensitivity of the AADS device may be possible by choosing a readout molecule with a high extinction coefficient (e.g. gold nanoparticles [E. J. Medcalf, E. Schäfer, F. Hollfelder, in preparation]). For example, Probst *et al*.^16^ used nanoparticles, a combination of confocal optical systems and a post-processing algorithm to achieve a sensitivity of 800 nM. Broadband-enhanced cavity absorption with mirrors could be added^27,28^, although manufacturing is complex due to needing to precisely place micromirrors on either side of the channel between the optical fibres.

On the other hand, AADS is cheaper to build (requiring no lenses and expensive laser equipment) than FADS, can be used on a benchtop (without laser protection) and provides a very accessible set-up for ultrahigh throughput screening. The robustness of absorbance sorting is enhanced by the practical measures of refractive index matching and baseline offsetting that will make it easier to obtain good quality quantitative data, and an increased sorting frequency ensures accurate sorting decisions based on high quality peak detection at lower volumes and high frequency. Post-processing analysis of droplet data can additionally be easily reviewed through the Python scripts provided, and has been written with packages that are consistently improved by the community, for easy updating of the code. Also, since the sorting uses Arduino-like microcontrollers, improvement in hardware will result in increased computation speed whilst retaining the accessibility of being inexpensive and open-source. Complementary work by Richter et al.^29^ has shown increased sorting throughput and removal of droplet trace artifacts by using a combination of surface acoustic waves and micro-lenses in the form of an optical air cavity. We envision that a further upgrade to AADS designs could incorporate design improvements from both studies.

By making device designs (deposited on our repository DropBase https://openwetware.org/wiki/Dropbase:_UHT-AADS_Sorter) as CAD files) and new software described in the SI deposited on GitHub (https://github.com/fhlab/UHT-AADS) immediately available as open-source material we hope to facilitate uptake of ultrahigh throughput screening in droplets across the community and make it the method of choice for protein engineering by directed evolution.

**Table 2:** Summary of the improvements of the UHT-AADS compared to the previous designs (Gielen *et al*.^14^, Zurek *et al*.^17^).

| **UHT-AADS Addition compared to AADS** | **Improvement** |
| --- | --- |
| Faster sorting speed | Combination of points below to achieve sorting at 1000 Hz with close to 100% efficiency, a 10-fold increase of throughput. |
| RI-matching oil | Removes scattering signal from droplet edges in the recorded traces and allows detection of droplets with a lower concentration of substrate and reduced volume. |
| Single layered inlet | Allows separation of smaller droplets individually and to droplets be evenly spaced. |
| Bias oil | Acts as a barrier to droplets from entering the positive outlet without dielectrophoresis. Allows transfer of positive hits to oil and surfactants solution dedicated for storage of stable emulsion |
| Faster Sorting algorithm with gating | Sorting algorithm capable of sorting at 4kHz with an Arduino Due. Adds sorting gates to select for specific residence time and voltage ranges. |

## Acknowledgements

This work was supported by a studentship from the BBSRC Doctoral Training Account, (E.J.M., BB/M011194/1), a Trinity College / Benn W Levy SBS DTP PhD Studentship (M.G.), a H2020 Marie Curie Fellowship by the European Union (T.S.K., 750772); the H2020 project MetaFluidics (F.H., 685474) and an H2020 ERC Advanced Investigator Award (F.H., 695669). We thank Esther Richter, Raymond Sparrow, and Thomas Franke for their input and helpful discussion of the manuscript, and Stefan Heimersheim for help with statistics.

## References

1. Obexer, R. et al. Emergence of a catalytic tetrad during evolution of a highly active artificial aldolase. Nat Chem 9, 50–56 (2017).

2. Fernández, D. S., Klein, O. J., Kaminski, T. S., Colin, P.-Y. & Hollfelder, F. Ultrahigh-throughput directed evolution of a metal-free α/β-hydrolase with a Cys-His-Asp triad into an efficient phosphotriesterase. Biorxiv 2022.02.14.480337 (2022) doi:10.1101/2022.02.14.480337.

3. Debon, A. et al. Ultrahigh-throughput screening enables efficient single-round oxidase remodelling. Nat Catal 2, 740–747 (2019).

4. Ma, F. et al. Efficient molecular evolution to generate enantioselective enzymes using a dual-channel microfluidic droplet screening platform. Nat Commun 9, 1030 (2018).

5. Gielen, F., Colin, P.-Y., Mair, P. & Hollfelder, F. Protein Engineering, Methods and Protocols. Methods Mol Biology Clifton N J 1685, 297–309 (2017).

6. Colin, P. et al. Ultrahigh-throughput discovery of promiscuous enzymes by picodroplet functional metagenomics. Nat Commun 6, 10008 (2015).

7. Neun, S. et al. Functional metagenomic screening identifies an unexpected β-glucuronidase. Nat Chem Biol 1–8 (2022) doi:10.1038/s41589-022-01071-x.

8. Tauzin, A. S. et al. Investigating host-microbiome interactions by droplet based microfluidics. Microbiome 8, 141 (2020).

9. Mahler, L. et al. Highly parallelized droplet cultivation and prioritization of antibiotic producers from natural microbial communities. Elife 10, e64774 (2021).

10. Scheele, R. A. et al. Droplet-based screening of phosphate transfer catalysis reveals how epistasis shapes MAP kinase interactions with substrates. Nat Commun 13, 844 (2022).

11. Matuła, K., Rivello, F. & Huck, W. T. S. Single-Cell Analysis Using Droplet Microfluidics. Adv Biosyst 4, 1900188 (2020).

12. Neun, S., Zurek, P. J., Kaminski, T. S. & Hollfelder, F. Ultrahigh throughput screening for enzyme function in droplets. Methods Enzymol 643, 317–343 (2020).

13. Neun, S., Kaminski, T. S. & Hollfelder, F. Methods in Enzymology. Methods Enzymol 628, 95–112 (2019).

14. Gielen, F. et al. Ultrahigh-throughput–directed enzyme evolution by absorbance-activated droplet sorting (AADS). Proc National Acad Sci 113, E7383–E7389 (2016).

15. Duncombe, T. A., Ponti, A., Seebeck, F. P. & Dittrich, P. S. UV–Vis Spectra-Activated Droplet Sorting for Label-Free Chemical Identification and Collection of Droplets. Anal Chem 93, 13008–13013 (2021).

16. Probst, J., Howes, P., Arosio, P., Stavrakis, S. & deMello, A. Broad-Band Spectrum, High-Sensitivity Absorbance Spectroscopy in Picoliter Volumes. Anal Chem 93, 7673–7681 (2021).

17. Zurek, P. J., Hours, R., Schell, U., Pushpanath, A. & Hollfelder, F. Growth amplification in ultrahigh-throughput microdroplet screening increases sensitivity of clonal enzyme assays and minimizes phenotypic variation. Lab Chip 21, 163–173 (2020).

18. Mao, Z. et al. Label-Free Measurements of Reaction Kinetics Using a Droplet-Based Optofluidic Device. J Laboratory Automation 20, 17–24 (2014).

19. Zurek, P. J., Knyphausen, P., Neufeld, K., Pushpanath, A. & Hollfelder, F. UMI-linked consensus sequencing enables phylogenetic analysis of directed evolution. Nat Commun 11, 6023 (2020).

20. Zachos, I. et al. Hot Flows: Evolving an Archaeal Glucose Dehydrogenase for Ultrastable Carba-NADP+ Using Microfluidics at Elevated Temperatures. Acs Catal 12, 1841–1846 (2022).

21. Sciambi, A. & Abate, A. R. Accurate microfluidic sorting of droplets at 30 kHz. Lab Chip 15, 47–51 (2014).

22. Baret, J.-C. et al. Fluorescence-activated droplet sorting (FADS): efficient microfluidic cell sorting based on enzymatic activity. Lab Chip 9, 1850 (2009).

23. Zinchenko, A. et al. One in a Million: Flow Cytometric Sorting of Single Cell-Lysate Assays in Monodisperse Picolitre Double Emulsion Droplets for Directed Evolution. Anal Chem 86, 2526–2533 (2014).

24. Virtanen, P. et al. SciPy 1.0: fundamental algorithms for scientific computing in Python. Nat Methods 17, 261–272 (2020).

25. García, G. et al. A Survey on FPGA-Based Sensor Systems: Towards Intelligent and Reconfigurable Low-Power Sensors for Computer Vision, Control and Signal Processing. Sensors 14, 6247–6278 (2014).

26. Salmon, A. R. et al. Monitoring Early-Stage Nanoparticle Assembly in Microdroplets by Optical Spectroscopy and SERS. Small 12, 1788–1796 (2016).

27. Rushworth, C. M., Jones, G., Fischlechner, M., Walton, E. & Morgan, H. On-chip cavity-enhanced absorption spectroscopy using a white light-emitting diode and polymer mirrors. Lab Chip 15, 711–717 (2014).

28. Neil, S. R. T., Rushworth, C. M., Vallance, C. & Mackenzie, S. R. Broadband cavity-enhanced absorption spectroscopy for real time, in situ spectral analysis of microfluidic droplets. Lab Chip 11, 3953–3955 (2011).

29. Richter, E. S. et al. Sorting of droplets at kHz rates using absorbance activated acoustic sorting. bioRxiv (2022).

