## Supporting Information for "Ultrahigh-throughput Absorbance Activated Droplet Sorting (UHT-AADS) for enzyme screening at kilohertz frequencies"

### Table of Contents

1. Detailed microfabrication protocol S3
2. Droplet sorting algorithm. S6
3. Droplet detection post-processing script and graph generator (no edges). S8
4. Droplet detection post-processing script and graph generator (edges). S11
5. Figure S1. Mean absorbance at 340 nm for enrichment experiment. S13
6. Supporting Information References. S14

### 1. Detailed microfabrication protocol

**1.1. Photolithography of microfluidic moulds**

The channel layout for the microfluidic chips was designed using AutoCAD (Autodesk) and printed out on a high-resolution film photomask (Micro Lithography Services). The designs in Fig. 1 are deposited on https://openwetware.org/wiki/Dropbase:_UHT-AADS_Sorter. The microfluidic devices were fabricated following standard hard lithography protocols that can be performed in the local cleanrooms or outsourced to contract manufacturing companies. First, microfluidic moulds were patterned on 3” size silicon wafer (Microchemicals) using high-resolution film masks (Microlithography Services Ltd) and SU-8 2050 photoresists (Kayaku Advanced Materials). A MJB4 mask aligner (SÜSS MicroTec) was used to UV expose all the SU-8 spin-coated wafers. The thickness of the structures (corresponding to the depth of channels in the final microfluidic devices) was measured using a DektakXT Stylus profilometer (Bruker).

We used the following settings for photolithography:

| | **Fabrication step (no. of layer)** | |
|---|---|---|
| | **1st layer** | **2nd layer** |
| Nominal thickness | 50 µm | 50 µm 2nd layer (100 µm final thickness) |
| Resist used | SU-8 2050 | SU-8 2050 |
| Spin coating speed | 1st step: 10 sec, 500 rpm<br>2nd step: 30 sec, 3500 rpm | 1st step: 10 sec, 500 rpm<br>2nd step: 30 sec, 4300 rpm |
| Pre-baking | 3 min at 65°C<br>6 min at 95°C | 3 min at 65°C<br>6 min at 95°C |
| Exposure (at ~10 mW $cm^2$) | 2 x 8.5 sec | 2 x 8.5 sec |
| Post-baking | 1 min at 65°C<br>6 min at 95°C | 1 min at 65°C<br>6 min at 95°C |
| Development in the beaker filled with 30-50mL of PGMEA (propylene glycol methyl ether acetate, Sigma-Aldrich) | Approximately 3-5 minutes until all uncured SU-8 is removed from the wafer; development time depends on the intensity of manual agitation. The development step after the 1st deposition is performed only for a 1-layer chip. | Approximately 5-10 minutes until all uncured SU-8 is removed from the wafer, development time depends on the intensity of manual agitation. |

| | | |
|---|---|---|
| Hard baking (optional) | 5 min at 150 °C (only for a 1-layer chip) | 5 min at 150 °C |
| Measured range of thicknesses | 46-47 µm | 96-97 µm |

**1.2 Soft lithography**

To manufacture PDMS microfluidic devices, 20-30 grams of silicone elastomer base and curing agent (Sylgard 184, Dow Corning) were mixed at a 10:1 (w/w) ratio in a plastic cup and degassed in a vacuum chamber for 30 minutes. PDMS was then poured on a SU-8 master wafer with SU-8 structures and cured in the oven at 65°C for at least 4 hours. Next, the inlet holes were punched using 1.0 mm diameter biopsy punchers with plungers (Kai MedicalIndustries). The patterned PDMS chip was then plasma bonded first to approximately 1-mm-thin PDMS slab (cured beforehand) and then to a 52 mm x 76 mm x 1 mm (length x width x thickness) glass slide (VWR) in a low-pressure oxygen plasma generator (Femto, Diener Electronics). As a result, we obtained a 3-layer device with patterned PDMS on top, thin PDMS slab in the middle, and the glass slide at the bottom. Next, the hydrophobic modification of microfluidic channels was performed by flushing the device with 1% (v/v) trichloro(1H,1H,2H,2H-perfluorooctyl)-silane (Sigma-Aldrich) in HFE-7500 (3M) and baked on a hot plate at 75 °C for at least 30 minutes to evaporate the fluorocarbon oil and silane mix.

**1.3 Integration of optical fibres with the sorting chip**

The fabrication of the sorting chip required additional integration with incident light and detection multimode optical fibres with SMA connector type, cladding diameter of 125 µm and a core diameter of 50 µm with numerical aperture (NA) of 0.22. A single 5m-long fibre optic patch cable (cat. no M14L05, Thorlabs) was cut at the middle and the outer protective PVC jacket was removed using a three-hole fibre stripper (cat. no FTS4, Thorlabs). Next, the Kevlar protective threads were cut with a scalpel and finally acrylate coating was removed using a fibre stripping tool (cat. no T06S13, Thorlabs). In the next step the tip of the fibre tip was cleaved using a ceramic fibre scribe (cat. no CSW12-5, Thorlabs) in order to obtain a flat tip end. The quality of the cleavage was inspected by passing a low power light through the fibre and a visual inspection of the shape of a beam emerging from the fibre tip end. If necessary, the cleavage was repeated until the spherical shape of the light beam was observed. The fibre ends were fixed to the microfluidic chip at least 1 day prior to an experiment. The fibre fixing process was performed on the microscope stage and a microscope camera was used to verify the position of the fibre ends. First, the microfluidic channels housing the fibres were filled with liquid PDMS and then fibre tips were manually inserted to the chip. Fibres were additionally stabilised by attaching them to the glass slide of the chip with epoxy glue (Araldite® Two Component Epoxy Paste Adhesive). The whole chip was left overnight on the microscope stage to let the PDMS and glue cure at room temperature. Such chips can be used for several sorting experiments, provided that they are washed carefully with pure HFE-7500 and dried by blowing with compressed air over them after each use.

### 2. Droplet sorting algorithm

The droplet detection algorithm for Arduino-like microcontrollers including gating for residence time and voltage at high frequencies. This allows for sorting using droplet size and the ability to exclude electrode activation for events such as air bubbles and dust. Additionally, ‘negative’ sorting can be achieved (selecting for a decrease in absorbance compared to the negative population) through selecting for a minimum voltage. The maximum sorting capabilities for an Arduino Due is around 4 kHz. The code is modified from Zurek *et al.*[1] and is written in the C programming language.

```
//Enter Values Here
const float sortThresh = 6; // in volts
const float dropletThresh = 2; // in volts
const float minVoltage = 4; // in voltz
const int minResidence = 2; // in microseconds
const int maxResidence=5000; // in microseconds
const int outputSource = 13; //connect this pin to function generator
const int inputSource = A0; //connect photodetector input to this pin

//Other Global Variables
float absRead;
long dropletWidth;
long dropletTimerEnd;
long dropletTimerStart;

// Absorbance read
float reading(){
absRead = analogRead(inputSource);
absRead = absRead/4096*10; //change 4096 depending on analogReadResolution
return absRead;
}

void setup() {
  Serial.begin(9600);
  pinMode(outputSource, OUTPUT);
  analogReadResolution(12);
 REG_ADC_MR = (REG_ADC_MR & 0xFFF0FFFF) | 0x00020000; //faster analog read
}

// Pulses according to parameters
void pulse(){
delayMicroseconds(50);
digitalWrite(outputSource, HIGH);
delayMicroseconds(50);
digitalWrite(outputSource, LOW);
}
```

```
// Advanced sorting algorithm
void sortAdv(){
 absRead = reading();
 if (absRead < dropletThresh){
  if (absRead < sortThresh){
   while (absRead < dropletThresh && absRead > minVoltage){
    absRead = reading();
   }
   dropletTimerEnd = micros();
   dropletWidth = dropletTimerEnd-dropletTimerStart;
   if (dropletWidth > minResidence && dropletWidth < maxResidence){
     pulse();
   }
  }
 }
 else {
  dropletTimerStart = micros();
 }
}

// Main loop
void loop(){
 sortAdv();
}
```

#### 3. Droplet detection post-processing script and graph generator (no edges)

The post-processing script for detection of droplet peaks and extraction of data including: number of droplets, frequency, baseline value, number of droplets above the threshold, standard deviation of droplets above the threshold, and gating data. Additionally graphs are produced with and without gating showing the raw data, a histogram plot of droplets and residence time against voltage. The scipy package was used to infer droplet data and is therefore compatible with future updates to this package. The code was written in the Python programming language.

```
import pandas as pd
import peakutils.baseline
from matplotlib import pyplot as plt
import matplotlib.gridspec as gridspec
import scipy.signal
import numpy as np


droplet_threshold = 9.9 #adjust to desired threshold
hits_threshold = 9.7 #adjust to desired threshold
distance_between_peaks = 20 # number of samples between neighbouring peaks
relative_height = 0.5 # the relative height at which peak width is measured, e.g. 0.5 is at half the prominence height
sample_rate = 263.000000/9994  #sample rate of detector
bins = 100 # number of histogram bins
residence_time_low = 0.1 #minimum residence time for gating in microseconds
residence_time_high = 0.3 #maximum residence time for gating in microseconds

file = "INSERT FILE" # file should be a csv with a “Time” column and a “Voltage” column with headers
df = pd.read_csv(file)
df = df[5000:6000] # select data range or comment out to use all data

#convert to pandas dataframe
Time = df["Time"]
Time = Time.to_numpy()
total_time = Time[-1]
total_time_s = total_time / 1000
print("Time interval measured:",np.round(total_time,decimals = 3), "milliseconds")
Voltage = df["Voltage"]
Voltage = Voltage.to_numpy()

#baseline detection
baseline = peakutils.baseline(-Voltage)
average_baseline = -np.mean(baseline)
```

```
print("Baseline at:", np.round(average_baseline, decimals = 3))

#peak detection
minima_V = Voltage*-1
minima = scipy.signal.find_peaks(minima_V,distance=distance_between_peaks,height=-
droplet_threshold)
minima_sort =
scipy.signal.find_peaks(minima_V,distance=distance_between_peaks,height=(-
hits_threshold,-9.25))
points, _ = minima
points_sort, _ = minima_sort
min_pos = Time[minima[0]]
min_height = minima_V[minima[0]]
peak_widths = scipy.signal.peak_widths(minima_V,points, rel_height=relative_height)
widths = peak_widths[0]
residence_time = widths*sample_rate

#graphs without gating

gs = gridspec.GridSpec(2,2, wspace = 0.5, hspace = 0.5, top = 0.94)

#voltage against time
fig = plt.figure()
ax1 = fig.add_subplot(gs[1,:])
ax1.plot(Time,Voltage)
ax1.scatter(min_pos,min_height*-1,color='gold',s=50,marker='X')
plt.xlabel("Time [ms]")
plt.ylabel("Detection Signal [V]")
ax1.grid()

#voltage against residence time
ax2 = fig.add_subplot(gs[0,1])
plt.scatter(residence_time,Voltage[points], alpha=0.1)
plt.xlabel("Residence time [us]")
plt.ylabel("Voltage [V]")

#histogram of droplet detection signals [V]
ax3 = fig.add_subplot(gs[0,0])
plt.hist(min_height*-1,bins=bins)
plt.xlabel("Detection signal [V]")
plt.ylabel("Relative frequency")
fig.suptitle(file, fontsize = 8)
plt.show()

#printing of detected data
frequency = points.size/total_time_s
print(np.around(frequency,decimals = 1),"droplets per second")
print(points.size, "droplets detected")
```

```
print(points_sort.size,"droplets detected above threshold")
V_values_above_threshold = Voltage[points_sort[:99]]
print("Standard deviation of droplet voltages detected above
threshold:",np.round(np.std(V_values_above_threshold), decimals = 3), "volts")

#peak detection for gating and printing of detected data
peak_widths = scipy.signal.peak_widths(minima_V,points_sort, rel_height=relative_height)
widths = peak_widths[0]
residence_time = widths*sample_rate
residence_time_gated = residence_time > residence_time_low
residence_time_gated = residence_time[np.where((residence_time >= residence_time_low)
& (residence_time <= residence_time_high))]
points1 = points_sort[np.where((residence_time > residence_time_low) & (residence_time <
residence_time_high))]
gated_voltage = Voltage[points1]
print(len(residence_time_gated),"droplets detected with gate")

#graphs with gating

#gated histogram of droplet detection signals
gs = gridspec.GridSpec(1,2, wspace = 0.5, hspace = 0.5, top = 0.94)
fig = plt.figure()
ax4 = fig.add_subplot(gs[:,1])
plt.hist(gated_voltage,bins=bins)
plt.xlabel("Detection signal [V]")
plt.ylabel("Relative frequency")
plt.title("Voltage with Gated Residence Time")

#gated voltage against residence time
ax4 = fig.add_subplot(gs[0,0])
plt.scatter(residence_time_gated,gated_voltage, alpha=0.5)
plt.xlabel("Residence time [us]")
plt.ylabel("Voltage [V]")
plt.title("Gated Residence Time")
plt.show()
```

### 4. Droplet detection post-processing script and graph generator (edges)

The post-processing script for detection of droplet peaks and extraction of data as above, but for when peaks are masked by droplet edges.

```
import numpy as np
from scipy.signal import argrelextrema
import scipy.signal
import pandas as pd
import peakutils.baseline
from matplotlib import pyplot as plt

file = "Insert file"
df = pd.read_csv(file)
df = df[100000:101000]

Time = df["Time"]
Time = Time.to_numpy()
total_time = Time[-1]
total_time_s = total_time / 1000
print("Time interval measured:",np.round(total_time,decimals = 3), "milliseconds")
Voltage = df["Voltage"]
Voltage = Voltage.to_numpy()

baseline = peakutils.baseline(-Voltage)
average_baseline = -np.mean(baseline)
print("Baseline at:", np.round(average_baseline, decimals = 3))

threshold_max_higher_lower_bounds = 10.2
# for local maxima
local_max_higher = argrelextrema(Voltage, np.greater)[0]
local_max_higher =
local_max_higher[(Voltage[local_max_higher]>threshold_max_higher_lower_bounds)]
threshold_max_lower_upper_bound = 7
threshold_max_lower_lower_bounds = 2
# for local maxima
local_max_lower = argrelextrema(Voltage, np.greater)[0]
local_max_lower =
local_max_lower[(Voltage[local_max_lower]>threshold_max_lower_lower_bounds)]
local_max_lower =
local_max_lower[(Voltage[local_max_lower]<threshold_max_lower_upper_bound)]
# for local minima
local_min = argrelextrema(Voltage, np.less)[0]
local_min = local_min[(Voltage[local_min]<8)&(Voltage[local_min]>4)] #change numerical
values to match dataset
local_max = np.append(local_max_higher,local_max_lower)
```

```
fig = plt.figure(figsize=(8,4))

plt.plot(Time,Voltage)
plt.scatter(Time[local_min],Voltage[local_min],color='gold',s=50,marker='X')
plt.scatter(Time[local_max],Voltage[local_max],color='gold',s=50,marker='X')
fig.suptitle(file, fontsize = 8)
plt.autoscale(enable=True, axis='x', tight=True)
plt.show()
```

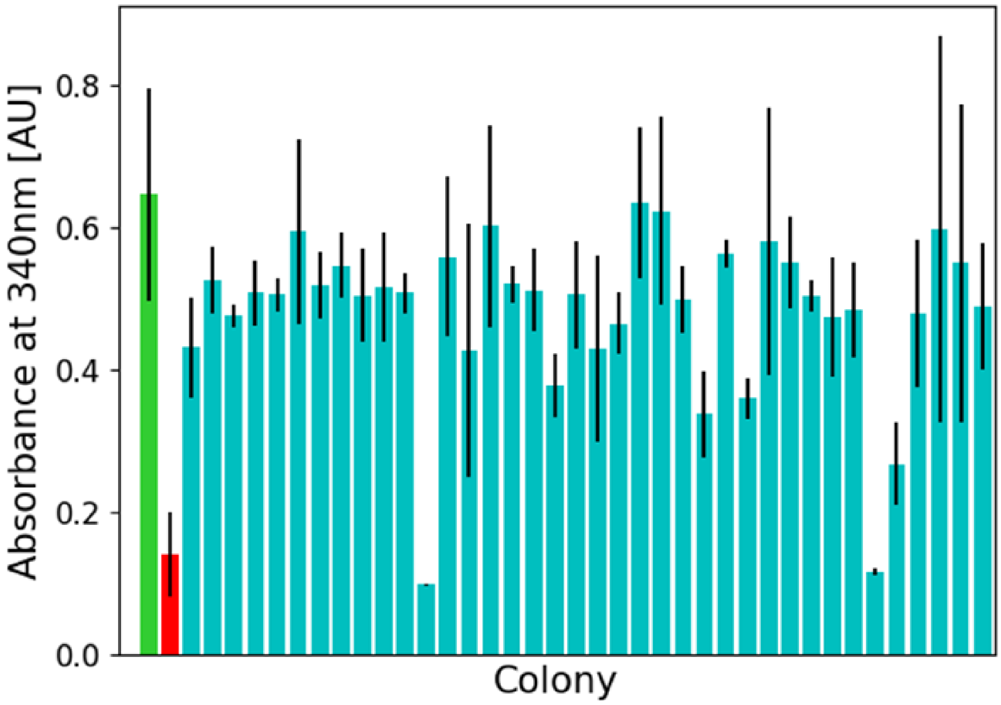


**Fig. S1.** Mean absorbance at 340 nm for 38 colonies (cyan, n=3 for each) picked after transformation of DNA collected from the positive outlet of the AADS device sorting at one kilohertz. The green bar shows the mean (n=9) of the positive control (pASK expressing wild type PheDH) and the red bar shows the mean (n=9) of the negative control (pASK expressing glycosidase). Error bars represent the standard deviation.

### 6. Supporting Information References

1. Zurek, P. J., Hours, R., Schell, U., Pushpanath, A. & Hollfelder, F. Growth amplification in ultrahigh-throughput microdroplet screening increases sensitivity of clonal enzyme assays and minimizes phenotypic variation. *Lab Chip* 21, 163–173 (2020).
